## Supplementary figures and images for "Anti-CD19 CAR T cells potently redirected to kill solid tumor cells"

### Supplemental Fig 1

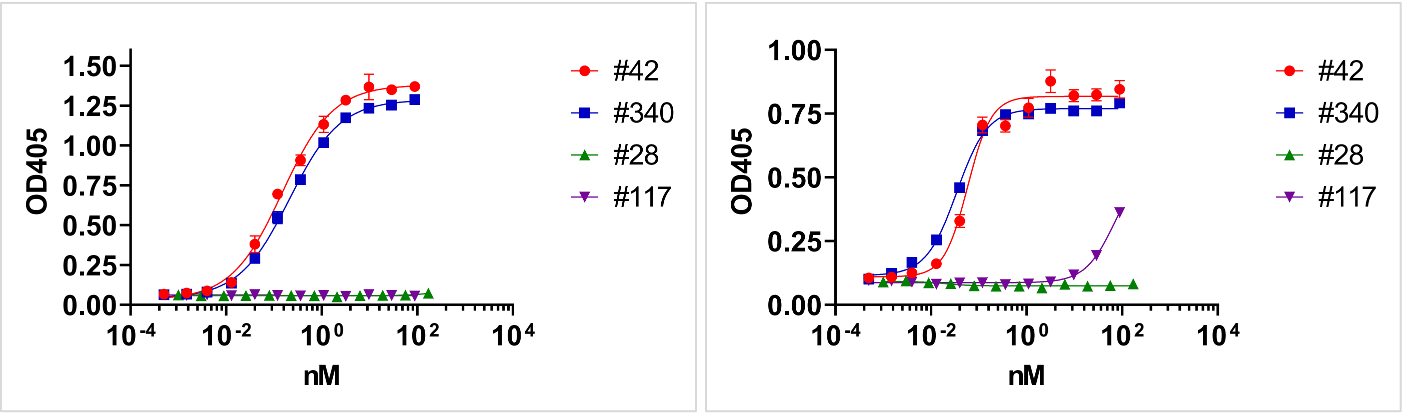
